# Male boldness and female parental care predict reproductive success in a bi-parental cichlid, the rainbow krib (*Pelvicachromis pulcher*)

**DOI:** 10.1101/2023.11.23.568478

**Authors:** U. Scherer, P.T. Niemelä, W. Schuett

## Abstract

1. Animal personality differences, i.e. consistent among-individual behavioural differences within populations, are prevalent across the animal kingdom. However, we are just beginning to understand the adaptive significance of the observed behavioural variation. We are particularly in need of empirical studies testing hypotheses of proposed theoretical frameworks aiming to understand the existence animal personality differences. In this study, we investigated a hypothesis derived from a framework suggesting that sexual selection may generate and maintain personality variation. The authors of this framework propose parental care as a mechanism linking animal personality and reproductive fitness.
2. We repeatedly measured individual boldness in male and female rainbow kribs, *Pelvicachromis pulcher*, a West African cichlid known for territorial cave breeding and shared parental care. We then formed 54 breeding pairs with varying behavioural contrasts in boldness. For pairs that produced offspring (N = 20), we repeatedly recorded parental care behaviour (parental boldness and brood guarding) of both parents over one month. Reproductive success was measured as the likelihood to reproduce, the number of offspring, and offspring size at the end of breeding.
3. In both sexes, we found consistent among-individual differences in boldness before breeding and in parental care behaviours. Bolder males were less likely to reproduce but, when breeding was successful, produced bigger broods compared to less bold males. Female parental boldness positively correlated with the number of offspring produced. However, individual boldness before breeding did not predict parental care behaviour in either sex and we found no effects of the pairs’ contrast in prebreeding boldness on parental care coordination or reproductive success.
4. The outcomes of our study may suggest that among-individual variation in male boldness is maintained by males with different behavioural types playing different reproductive strategies of equal average fitness. Future research should delve into understanding the intricate link between male boldness and reproductive success, exploring for instance underlying genetic mechanisms and interactions with environmental conditions.

## INTRODUCTION

The existence of animal personality differences (consistent among-individual differences in behaviour within a population) is well documented throughout the animal kingdom (Bell et al., 2009; Gosling, 2001; Kralj-Fišer & Schuett, 2014; Stamps, 2007). Yet, we are just starting to understand the adaptive significance of the phenomenon (Moiron et al., 2020). Several theoretical frameworks aiming to uncover the evolutionary background of individual differences in behaviour have been proposed; these include, for instance, social advantages of behavioural consistency (Dall et al., 2004; McNamara et al., 2009; Wolf & McNamara, 2012), negative frequency-dependent (Wolf & McNamara, 2012) and fluctuating selection (Dingemanse et al., 2004; Dingemanse & Réale, 2005), state-dependent feedback loops and life-history trade-offs (Dammhahn et al., 2018; Ehlman et al., 2022; McNamara & Houston, 1996; Schuett et al., 2015), and sexual selection (Schuett et al., 2010). However, empirical studies that test the predictions of the above frameworks are still scarce.

Here, we focus on a hypothesis derived from the framework that sexual selection may generate and maintain personality variation (Schuett et al., 2010). It is proposed that, in caregiving species, certain behavioural traits expressed outside breeding are sexually selected for via mate choice because they predict future parental care behaviour (in one or both sexes). A non-breeding behaviour could be sexually selected for, if the sex choosing a partner obtains a direct fitness benefit from the partner’s behaviour per se. An example for a correlation between non-breeding behaviour and parental care quality is found in blue tits, *Cyanistes caeruleus*, where fast exploring females feed their chicks at higher rates and have higher reproductive success than slow exploring females (Mutzel et al., 2013). In this specific scenario, the framework would predict males to prefer highly exploratory females. Alternatively, certain behavioural type combinations within a pair may lead to higher reproductive success. This could be the case if within-pair behavioural compatibility is important for determining reproductive success through better cooperation and coordination of parental behaviours (Royle et al., 2010). For example, in long-tailed tits, *Aegithalos caudatus*, synchronised offspring provisioning behaviour increases reproductive success through a lower risk of predation (Bebbington & Hatchwell, 2016). Depending on the species’ biology and environmental conditions encountered, either positive or negative behavioural assortment of partners might lead to higher compatibility (Royle et al., 2010; Scherer, Kuhnhardt, et al., 2017; Schuett et al., 2010). While several studies show that consistent among-individual differences in behaviour matter during mate choice (Bierbach et al., 2021; Scherer, Kuhnhardt, et al., 2017; Schuett, Godin, et al., 2011), the potential role of parental care as the mechanism linking individual variation in behaviours expressed before breeding to variation in reproductive success has rarely been addressed (but see Chira, 2014; Schuett et al., 2011).

Here, we present a breeding experiment, in which we tested whether consistent among-individual variation in boldness expressed before breeding relates to consistent among-individual variation in parental care and reproductive success in the rainbow krib, *Pelvicachromis pulcher*, a West African cichlid. Rainbow kribs are territorial cave breeders and both parents care for their fry for several weeks, as can be observed in their sister species *P. taeniatus* (Thünken et al., 2010). In cichlids, breeding pairs often perform various parental activities and while each partner can carry out all tasks if necessary (Itzkowitz et al., 2001, 2005), partners seem to divide roles with the female typically providing more direct care (e.g. keeping the brood together, guiding them to feeding grounds, cleaning eggs) and the male engaging more in indirect care (e.g. defending or protecting offspring and eggs from con-and heterospecific intruders, territory patrolling and general vigilance) (Lavery & Reebs, 2010; McKaye & Murry, 2008; Richter et al., 2005). We tested male and female rainbow kribs for their boldness repeatedly and we then created breeding pairs that varied in how similar they were regarding their pre-breeding boldness. During breeding, we recorded indirect (parental boldness) and direct (brood guarding) parental care behaviour for both parents, and at the end of breeding we assessed reproductive success as the likelihood to reproduce, the number and size of offspring produced.

We expected (I) consistent among-individual variation in boldness expressed before breeding to predict reproductive success, and we predicted that (II) pre-breeding behavioural type and reproductive success are linked via parental care, i.e. we expected individual differences in pre-breeding boldness to predict differences in parental care behaviour, and in turn, individual differences in parental care to predict reproductive success. (I-II) We predict reproductive benefits may arise from behavioural dis-assortment because of the species’ reproductive system and female mating preference for males of a dissimilar level of boldness to themselves (Scherer, Kuhnhardt, et al., 2017), i.e., we expect dis-assortment to be associated with increased reproductive success via facilitation of the specialisation into distinct roles (direct vs. indirect care), reducing stress and conflict and easing the efficient coordination of care (Royle et al., 2010; Schuett et al., 2010). Alternatively, one might expect directional effects of male and/or female behavioural type on their parental care and reproductive success. As a basic assumption underlying our investigation, we expected among-individual variation in pre-breeding boldness as well as parental care behaviours.

## METHODS

### Experimental procedures

#### Study animals and housing conditions

Test fish were obtained from a house breed at the Universität Hamburg and local suppliers; and were maintained in same-sex groups matched for family and origin (100-200l tanks), at 25±1°C, and a 12:12 hours light:dark period. Tanks were endowed with a layer of sand (approx. 1 cm thick), an internal filter and plastic plants. Water changes (50%) were done once a week and fish were fed daily with live *Artemia* spp.. Four days before boldness tests, individuals were transferred to individual housing tanks (25L, 50 x 50 x 25 cm, same holding conditions as above) and measured for their standard length using ImageJ (Schneider et al. 2012) (mean ± SE standard length, males: 5.42 ± 0.05 cm, females: 4.39 ± 0.04 cm). All test fish were uniquely VIE-tagged for identification (Schuett et al., 2017).

#### Experimental Outline

We measured boldness before breeding as activity in the presence of a predator (see ‘Boldness tests’). Prior to the experiment, we validated the method: rainbow kribs decreased their activity when presented with an animated photo of a predator (*Parachanna obscura*, a naturally sympatric occurring predato*r*) compared to a control without the animated predator (a white computer screen), while there was no difference to the response towards the live predator (Scherer, Godin, et al., 2017). We then created breeding pairs that varied regarding their similarity in average boldness (see ‘Pairing and breeding’) and during breeding, we performed six parental care tests to assess parental boldness and brood guarding for each successfully reproducing breeding pair (see ‘Parental care tests’). We measured reproductive success as the likelihood to reproduce and the number and size of offspring produced after one month of care (see ‘Pairing and breeding’ for details).

#### Boldness tests

Before breeding, all males (N = 54) and females (N = 54) were tested for their boldness twice with five days in between trials. Boldness tests were performed following Scherer, Godin, et al. (2017). In short, individual test fish were exposed to an animated predator specimen (*P. obscura*), which was presented on a computer screen for 11 min. A photograph of the predator was animated to swim back and forth in front of a white background (see Scherer, Godin, et al., (2017) for details on the animation production). We assessed individual activity in the presence of the predator as the total number of squares visited (including revisits) for 10 min (no tracking of the first minute) using the tracking software Ethovision XT 11 (Noldus, Wageningen, The Netherlands). Therefore, test tanks (25×50 cm) were divided into 8 squares each measuring 12×12 cm squares. For all trials, we used a predator specimen the test fish had not seen before (N_predators_ = 4, mean ± SE standard length = 19.3 ± 0.3 cm). To create breeding pairs and for analyses (not repeatabilities), we used the average boldness an individual showed over both boldness tests (mean ± SE number of squares visited: males = 63.56 ± 4.26, females = 41.07 ± 3.64).

#### Pairing & breeding

Four days following the boldness tests, we set up breeding pairs (N = 54) of varying behavioural contrasts in boldness. That is, fish were tested in 5 batches with 12 males and 12 females per batch (N total number of fish tested = 60 males and females, respectively). Boldness was quantified directly after trials allowing us to form pairs based on the behaviour shown. For each batch, pairs were formed in a way that we minimized the behavioural contrast for half of the pairs, whereas the other half was maximized for within-pair behavioural contrast (varying sex of the bolder individual). To initiate breeding, we introduced the respective male and female into a breeding tank (50×50 cm, water level = 25 cm), equipped with half a clay pot as breeding cave, a plastic plant, a layer of sand, and an internal heater (all in a standardised position). We monitored the breeding cave for eggs daily, using a small dentist mirror (diameter = 3 cm). Breeding pairs that did not successfully spawn within 28 days were transferred back to their home tanks and were not further used in this experiment. Breeding pairs that did produce fry were allowed to raise their brood for 33 days (spawning = day 1). During this breeding period, we assessed parental care behaviour as outlined below. Different to housing conditions, we increased the number of water changes to three times a week and reduced the amount of water being changed to approx. 30% during the breeding period (no water changes on parental care test days).

For breeding pairs that reproduced (N = 20 breeding pairs), we counted all offspring produced (mean ± SE = 68.7 ± 9.4) and measured their size (mean ± SE = 1.56 ± 0.03 cm, standard length measured from photos using ImageJ, Schneider et al. (2012)) at the end of the breeding period (day 33). For statistical analyses, we averaged offspring size per brood.

#### Parental care tests

During the breeding period, we quantified parental boldness and brood guarding in the presence of an animated intruder. We used three different intruder types: a conspecific male, a conspecific female, and a predator (conspecifics are brood predators in the species). Each intruder type was used twice, i.e., each breeding pair was tested for their parental care behaviour six times with three days elapsing between successive trials. For each breeding pair, we randomised the testing order for the first time each intruder type was used (first, second and third parental care test) and then repeated parental care tests in the same order (six tests in total, e.g. a breeding pair was tested in the following order: male intruder, female intruder, predator, male intruder, female intruder, predator). We started our parental care observations one day after the offspring became free-swimming, i.e. day 10 post spawning. Fertilised *P. pulcher* eggs take three days to develop into wrigglers (free embryos), which stay in the breeding cave for approximately another five days. On day 9 following fertilisation, offspring become free-swimming fry and leave the breeding cave to search for food.

To start a parental care test, we introduced a tablet (Surftab Theatre, 13.3” Full-HD-IPS display; Trekstar, Bensheim, Germany) on a standardised side of the breeding tank showing one of the three intruder types for 11 min (Scherer, Godin, et al., 2017). We video-recorded the breeding pair’s response and manually assessed the activity for each parent from the videos (duration of video analysis was 10 min, starting 1 min after the start of the video). Similar to the procedure in the above boldness tests, parental boldness was assessed as the total number of squares visited (including revisits). Therefore, the breeding tank was divided into 16 squares each measuring 12×12 cm squares using markings alongside the vertical tank walls. Further, male and female brood guarding was quantified from snapshots, i.e., over the 10 min test period, we scored every 30 sec (21 frames in total) whether a parent was within one fish length distance to the brood (approx. 6 cm, or half a square) or not. If so, this was scored as brood guarding (Thünken et al., 2010). Individual brood guarding was then calculated as the number of frames where the individual was attending the brood in relation to all frames analysed (resulting in values ranging from 0 to 1: 0 = the parent was not attending the brood at all, 1 = the parent was with the brood throughout). All videos were analysed by the same observer (US). At the time of video analysis, the observer was not aware of pre-breeding boldness scores.

Male intruder sizes were matched to the male’s standard length and, similarly, female intruder sizes were matched to the female’s standard length (size difference ≤ 2 mm). Predator sizes were as described in the boldness test. Test fish were presented with unfamiliar intruder specimen only. The intruder type (male conspecific, female conspecific, predator) did not affect parental care behaviour, regardless of focal individual personality (**Supplementary information** “1 Intruder type did not affect parental care behaviour”). Consequently, we averaged individual parental care behaviour over all six parental care tests for statistical analyses (not repeatabilities).

### Data analyses

#### General Details

All data analyses were done in R version 4.2.1 (R Core Team, 2022). We fit (generalized) linear mixed-effect models ((G)LMMs) using the *lme4*-package (Bates et al., 2015). Wherever appropriate, we included test fish ID and family as random terms in (G)LMMs. Model fit was visually ensured using residual-and q-q plots. We selected most parsimonious models in a stepwise backward removal of non-significant terms. In the main text, we report p-values and model estimates for significant terms only; a complete summary with estimates and p-values for all full models (containing all predictors) and final models (containing significant predictors only) is provided in **Supplementary Information** “2 Full and final model summaries”. Model summary tables (including marginal and conditional R^2^ following Nakagawa et al. (2017)) were built using the R package *sjPlot* (Lüdecke, 2022). For all analyses (excluding repeatabilities), we used average behaviours shown over all trials, i.e., for each individual, pre-breeding boldness was averaged over both trials and parental care behaviour (parental boldness and brood guarding) was averaged over all six trials.

Offspring size and number were negatively correlated with each other (LMM with average offspring size per brood as response, the number of offspring produced as predictor; intercept ± SE = 1.683 ± 0.040, slope ± SE = –0.002 ± 0.000, p = 0.003, marginal R^2^= 0.375, N = 20). And in both sexes, parental boldness and brood guarding were negatively correlated (LMM with parental boldness as response and brood guarding as predictor, sexes were tested in separate models; males: intercept ± SE = 139.667 ± 12.936, coefficient ± SE = –78.351 ± 16.425, p <0.001, marginal R^2^= 0.139, N = 120; females: intercept ± SE = 85.220 ± 9.240, coefficient = –30.025 ± 11.423, p = 0.010, marginal R^2^ = 0.056; N = 120 for each sex). However, only a small to moderate amount of variation can be explained by those correlations, we therefore did not consider our measures to be redundant (Dormann et al. 2013). Female size did neither affect the number of offspring produced (LMM with offspring number as response and female size as predictor; p = 0.391, N = 20) nor offspring size (LMM with offspring size as response and female size as predictor; p = 0.495, N = 20).

#### Behavioural repeatabilities

We tested for significant consistent among-individual variation in boldness before breeding (N for each sex = 108 observations from 54 individuals) and in parental behaviours (parental boldness and brood guarding, N for each sex = 120 observations from 20 individuals, respectively) via calculating repeatabilities (R) following Hertel et al. (2020). In short, we built LMMs with the behaviour of interest as response and test fish ID as well as family as random terms, no predictors were included. We calculated repeatabilities for males and females separately. R is considered significant when its 95% CI does not overlap with zero.

#### (I) Does behavioural type predict reproductive success?

We wanted to know if (a) boldness expressed before breeding predicts whether a breeding pair produces fry and for those breeding pairs that did reproduce, we wanted to know if (b) boldness before breeding predicts brood and offspring size. (a) To test for effects of behaviour per se on the likelihood to reproduce, we ran a GLMM with a binary response variable encoding whether a breeding pair successfully reproduced (yes or no; N = 54 breeding pairs) and both male and female average pre-breeding boldness as predictor variables. We further included the interaction term between male and female boldness to test if the combination of male-female behavioural type in boldness affects a breeding pair’s likelihood to reproduce. (b) To answer our second question, we ran two linear models (LMs) with either offspring number or average offspring size as response. As before, we included both male and female average pre-breeding boldness as predictors (testing for an effect of behaviour per se) as well as the interaction term between both variables (testing for an effect of behavioural compatibility) (N = 20 broods).

#### (II) Does parental care link behavioural type to reproductive success?

The aim of our second hypothesis was twofold: we wanted to know if (a) boldness expressed individually, and before breeding, predicts parental care behaviour, and if (b) parental care predicts reproductive success. (a) To test if pre-breeding behavioural type in boldness predicts behavioural type in parental boldness, we built an LMM with average parental boldness as response and average pre-breeding boldness as predictor. The model was run for the two sexes separately (N = 20 individuals for both models). To test if the combination of male and female behavioural types predicts their coordination of care (i.e., do dis-assortative pairs show more pronounced role division?), we calculated male-female behavioural contrast in both pre-breeding boldness as well as parental boldness as average male behaviour minus average female behaviour (i.e., positive values indicate that the male is bolder than the female and vice versa). We then fit an LMM with male-female contrast in parental boldness as response and male-female contrast in pre-breeding boldness as predictor (N = 20 breeding pairs). In the same fashion, we proceeded with brood guarding. That is, we built two LMMs (one for each sex) with average brood guarding as response and average pre-breeding boldness as predictor (N = 20 individuals for each model), and a third LMM with parental contrast in brood guarding (male minus female behaviour) as response and contrast in pre-breeding boldness as predictor (N = 20 breeding pairs).

(b) To test whether parental care predicts reproductive success via effects of behaviour per se vs. effects of compatibility, we used essentially the same model structure as in (Ib). That is, we built two LMs with either offspring number or offspring size as response and average male and female parental boldness as well as the interaction term between those two variables as predictors. To test for effects of brood guarding, we built another two LMs with either offspring number or size as response and average male and female brood guarding as well as their interaction term as predictors.

## RESULTS

### Repeatable behavioural variation before and during breeding

Male and female rainbow kribs showed consistent among-individual variation in all three behaviours observed, i.e., in boldness expressed before breeding (repeatability males = 0.631, CI [0.555, 0.707]; repeatability females = 0.563, CI [0.470, 0.638]), parental boldness (repeatability males = 0.381, CI [0.269, 0.505]; repeatability females = 0.114, CI [0.063, 0.185]), and brood guarding (repeatability males = 0.181, CI [0.103, 0.277]; repeatability females = 0.182, CI [0105, 0.267]) (see **Supplementary information** “3 Behavioural profiles of males and females used in the breeding experiment” for a visualization of behavioural profiles).

### (I) Behavioural type predicted reproductive success

Bolder males were less likely to reproduce compared to less bold males (odds ratio ± SE = 0.978 ± 0.012, p = 0.049, **Figure 1b**; **Supplementary Table S3**). However, of those males that did reproduce, bolder males sired more offspring (estimate ± SE = 0.800 ± 0.193, p = 0.001, **Figure 2b**; **Supplementary Table S4**). Male boldness did not predict offspring size (**Figure 2a**; **Supplementary Table S4**). Neither female pre-breeding boldness nor the combination of male and female pre-breeding boldness predicted a pair’s likelihood to reproduce (**Figure 1a**; **Supplementary Table S3**) or their reproductive success in terms of the number and size of offspring produced (**Figure 2a**; **Supplementary Table S4**).

**Figure 1.**
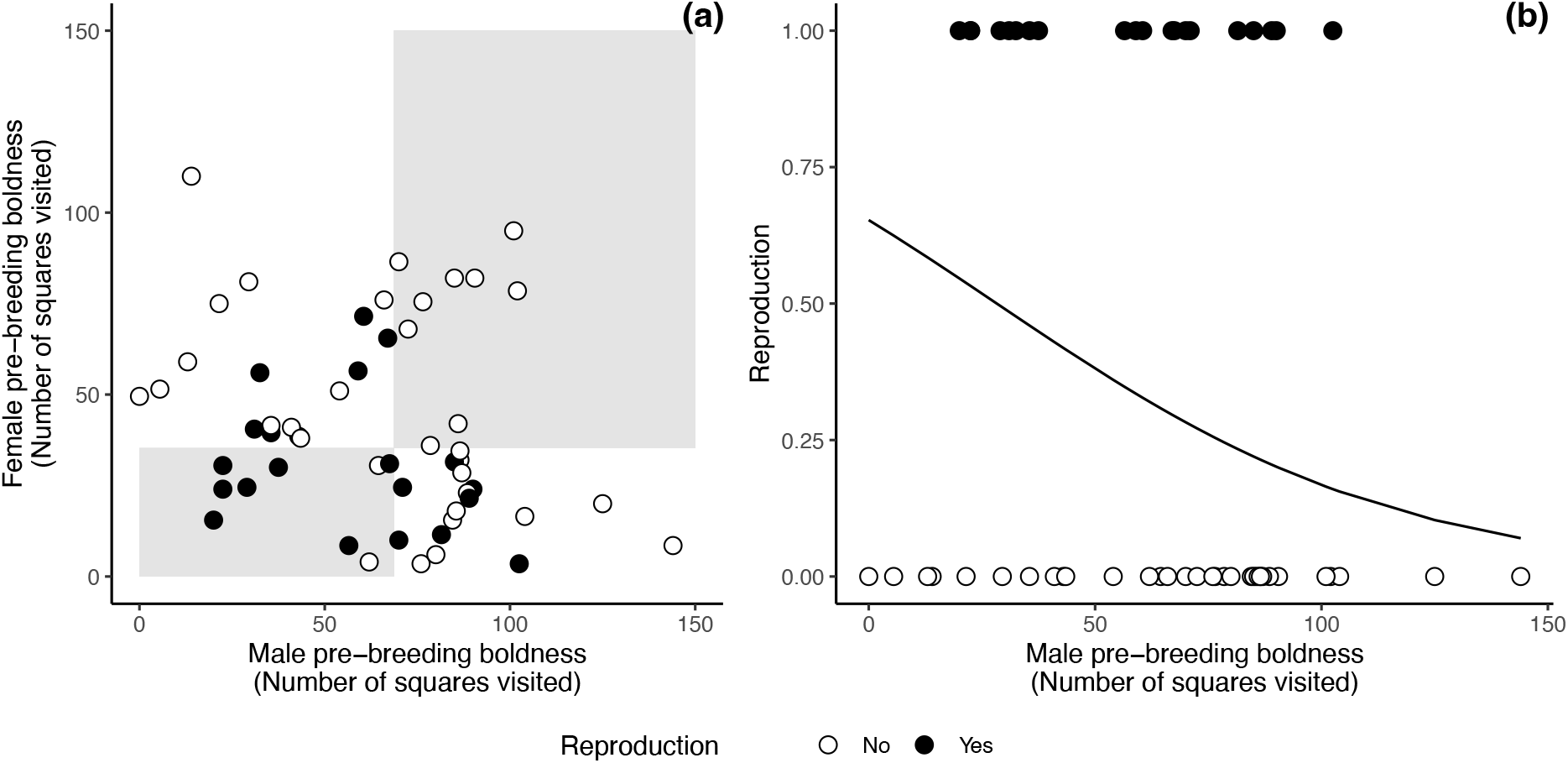
(a) The combination of male-female pre-breeding boldness did not predict a pair’s likelihood to reproduce but (b) bolder males were less likely to sire offspring. (a-b) Data points represent pairs that reproduced (black) and that did not reproduce (white). (a) X-and y-axes represent male and female boldness, respectively; their combined effect on the likelihood to reproduce reads as a pattern in space, i.e. a potential effect of behavioural assortment would be visible as an aggregation of either white or black data points in grey (assortative pairs where both individuals are either bold or less bold) vs. white (dis-assortative pairs with one bold and one less bold individual) areas of the coordinate system; bold vs. less bold threshold is the sex-specific median (for graphical purposes only). (a) The likelihood to reproduce in dependence of male boldness expressed before breeding is presented as a black regression line.

**Figure 2.**
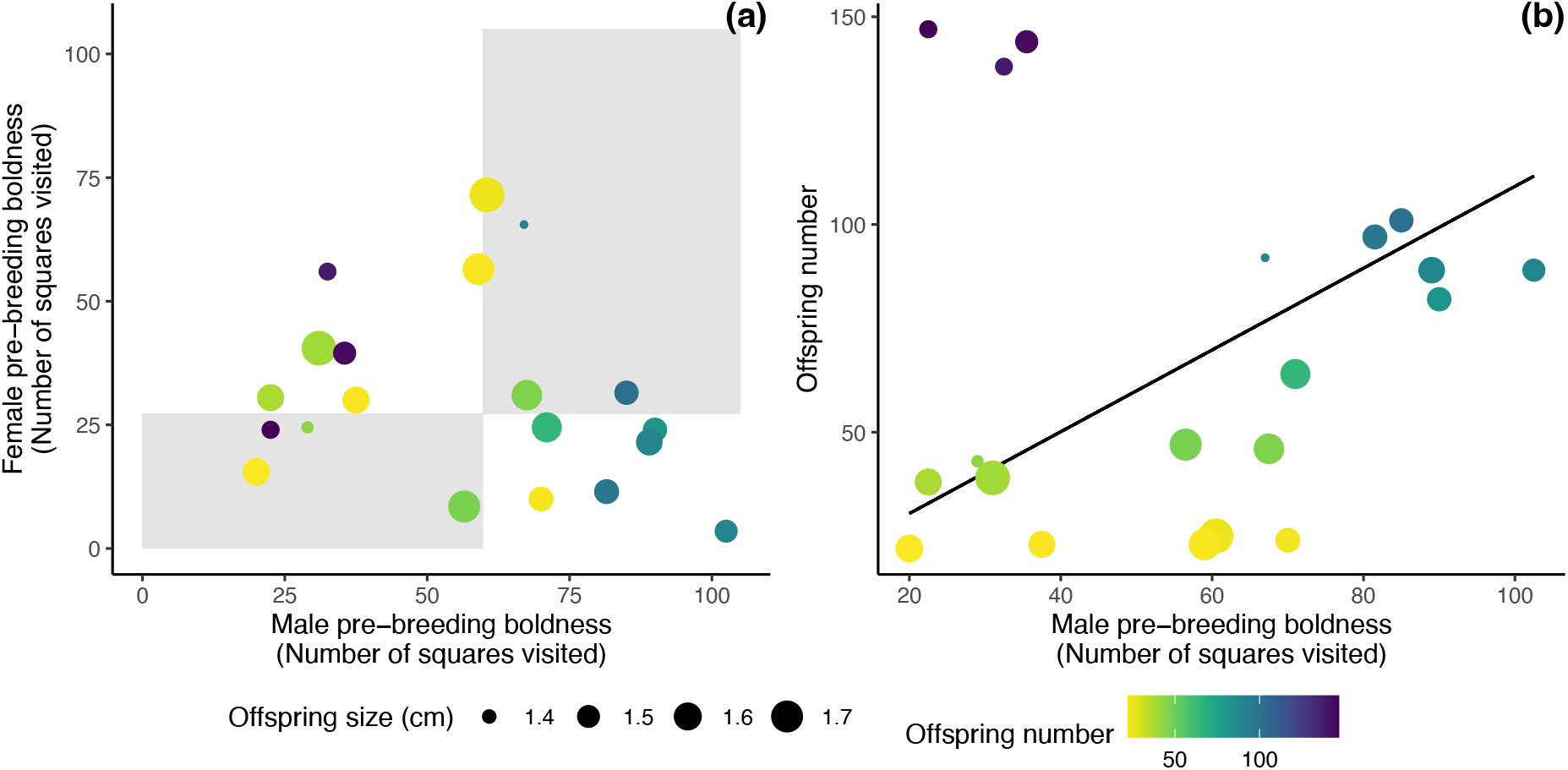
(a) The combination of male-female pre-breeding boldness did not predict a breeding pair’s reproductive success. On the graph, the combined effect male and female pre-breeding boldness on offspring size and number reads as potential pattern in space, i.e. an effect of behavioural assortment on offspring size/number would be visible as an aggregation of similarly sized/coloured data points in grey areas (assortative pairs where both individuals are either bold or less bold) vs. white areas (dis-assortative pairs with one bold and one less bold individual) of the coordinate system; bold vs. less bold threshold is the sex-specific median (for graphical purposes only). (b) Bolder males sired more offspring. (a-b) Data points represent the 20 broods that were produced in the breeding experiment; the colouration indicates brood size and the size of the data point indicates average offspring size for each brood.

### (IIa) Behavioural type did not predict parental care

There were no effects of behavioural type in pre-breeding boldness on expression of parental care, i.e. male and female pre-breeding boldness did not correlate with their parental boldness (**Figure 3a**, **c**; **Supplementary Table S5**) or brood guarding (**Figure 3b**, **d**; **Supplementary Table S6**). Furthermore, we found no effect of behavioural assortment on parental role division, i.e. male-female behavioural contrast in pre-breeding boldness did not predict their behavioural contrast in parental boldness (**Figure 3e**; **Supplementary Table S5**) or brood guarding (**Figure 3f**; **Supplementary Table S6**).

**Figure 3.**
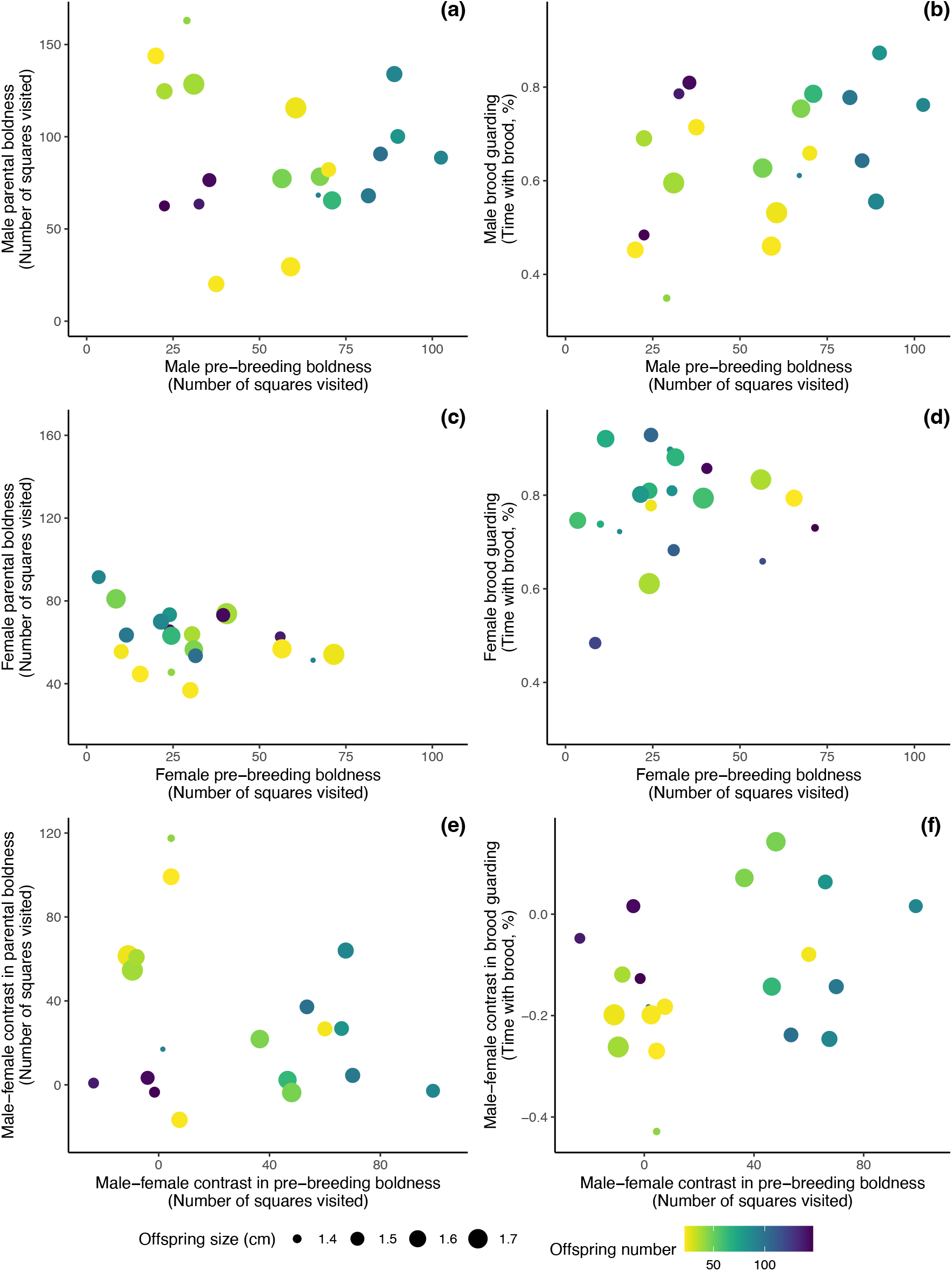
Male (a-b) and (c-d) female pre-breeding boldness did not predict their parental care behaviour (parental boldness and brood guarding). Likewise, (e-f) male-female contrast in pre-breeding boldness did not predict within-pair behavioural contrast in parental care. (a-f) Data points are coloured by brood size and the size of the data point indicates average offspring size for each brood (N = 20).

### (IIb) Link between maternal care and reproductive success

Female parental boldness was positively correlated with the number of offspring produced (estimate ± SE = 1.029 ± 0.465, p = 0.041, **Figure 4a** and **Figure 4c**; **Supplementary Table S7**). However, we found no connection between male parental boldness or the combination of male-female parental boldness on the number of offspring (**Figure 4a**; **Supplementary Table S7**), and likewise, there were no effects of parental care (male behaviour, female behaviour or male-female behavioural contrast in parental boldness or brood guarding) on offspring size (**Figure 4a**, **b**; **Supplementary Table S8**).

**Figure 4.**
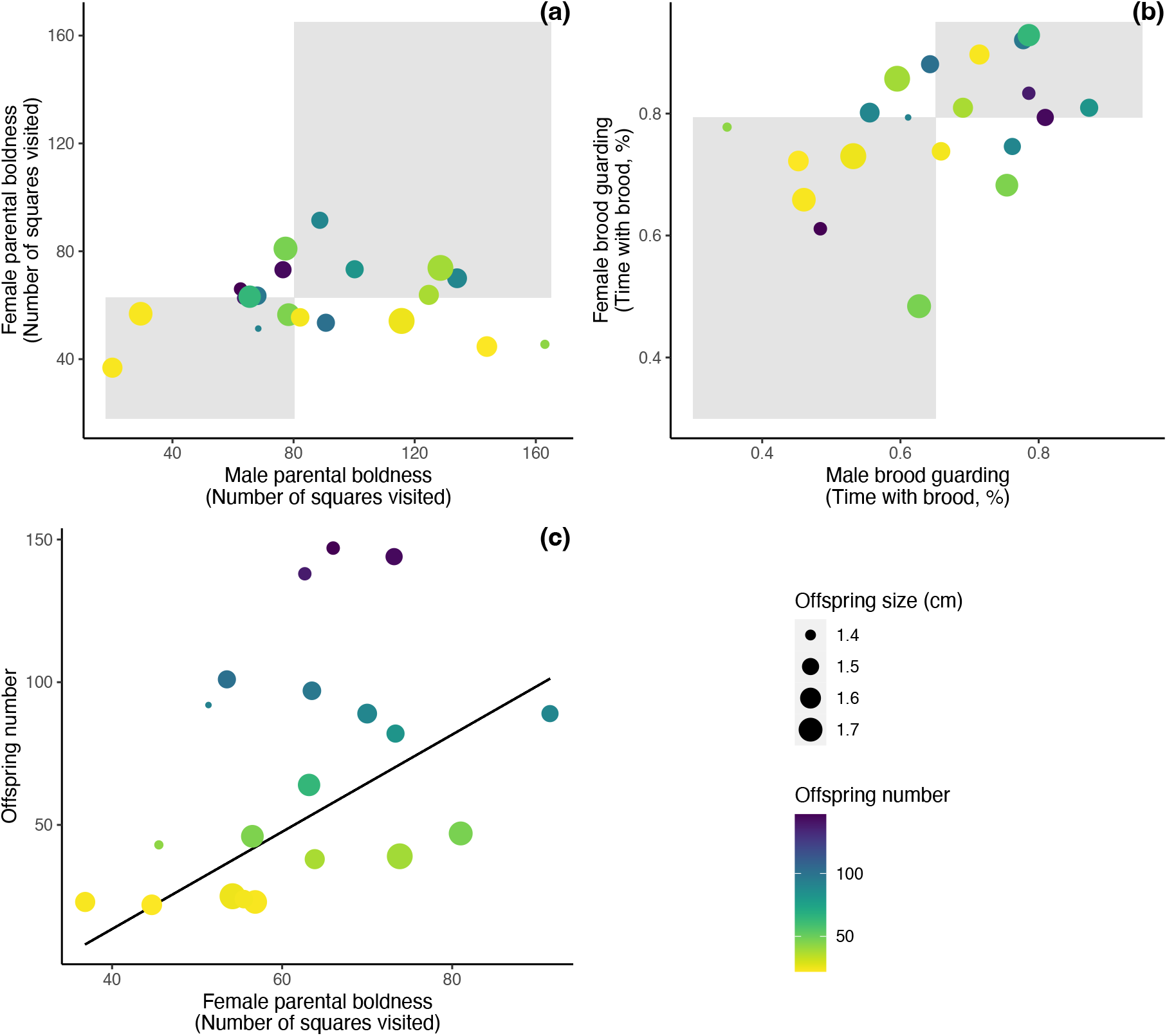
The combination of male-female (a) parental boldness or (b) brood guarding did not predict their reproductive success but (c) female parental boldness was positively correlated with the number of offspring produced. (a-c) Data points are coloured by brood size and the size of the data point indicates average offspring size per brood (N = 20). (a-b) X-and y-axes represent male and female behaviour, respectively; their combined effect on offspring size and number reads as potential pattern in space, i.e. an effect of behavioural assortment on offspring size/number would be visible as an aggregation of similarly sized/coloured data points in grey areas (assortative pairs where both individuals are either bold or less bold) vs. white areas (dis-assortative pairs with one bold and one less bold individual) of the coordinate system; bold vs. less bold threshold is the sex-specific median (for graphical purposes only).

## DISCUSSION

We found that male and female rainbow kribs showed consistent among-individual behavioural variation, i.e. animal personality differences, in boldness assessed before breeding as well as in both parental care behaviours (parental boldness and brood guarding). We found partial support for our first hypothesis (namely, individual differences in boldness predict reproductive success): bolder males were less likely to reproduce but sired more offspring when they did, however, female boldness did not predict reproductive success. Our results do not support our second hypothesis (among-individual variation in parental care links variation in pre-breeding boldness with variation in reproductive success): even though we found female parental boldness to be positively correlated with the number of offspring produced, pre-breeding boldness did not predict parental care behaviour in either sex.

Bolder males produced more offspring compared to less bold males. Since there were no direct (behavioural) benefits associated with male boldness expressed before breeding (i.e. no link to parental care behaviour), the reproductive benefit of bolder males (bigger broods) might be of indirect (genetic) nature, i.e. male intrinsic quality may cause males to be bolder as well as to sire more offspring. A recent meta-analysis showed that individuals expressing more risky behaviours (e.g. boldness) lived longer in the wild, indicating (at least indirectly) better quality of risky behavioural types (Moiron et al., 2020). A potential pathway linking male boldness and brood size in the present study could be that bolder males might have higher sperm quality (e.g. higher sperm count or velocity) and, therefore, higher insemination success. Empirical support for this hypothesis comes from the guppy (*Poecilia reticulata*): bolder guppies have higher sperm count (Gasparini et al., 2019), and they sire bigger broods (Herdegen-Radwan, 2019). Future studies may investigate a potential link between male boldness, sperm quality, and reproductive success in more detail.

Despite bolder males producing more offspring, they were less likely to reproduce at all compared to less bold males. The apparent contradiction may indicate that different male behavioural types follow different reproductive strategies. More specifically, because bold males sire more offspring per reproductive event, they might be able to afford to be choosier and to not realize as many mating opportunities. Males expressing lower boldness, on the other hand, may compensate for the lower output per reproductive event by realizing more mating opportunities. A recent meta-analysis supports this hypothesis by showing that attractive males are choosier than unattractive males (attractiveness being defined as traits that signal mate quality) (Dougherty, 2023). If the proposed reproductive strategies (bold males produce few but large broods and less bold males produce many but small broods) yield equal fitness outcomes, this could help maintain among-individual variation in male boldness. Upcoming work may address this prediction by quantifying reproductive success of males differing in their level of boldness under more natural settings (McCowan et al., 2014). It will be particularly informative to consider the potential role of environmental conditions like predation risk or mate availability on variation in reproductive success of different behavioural types. One might expect that, depending on the specific environment encountered, either a high or a low level of boldness might be associated with higher reproductive success, however, environmental heterogeneity over time and/or in space can lead to balancing selection and the maintenance of among-individual variation in male boldness (Dingemanse et al., 2004; Mangel, 1991; Penke et al., 2007).

The observed pattern of less bold males having a higher likelihood of reproduction could also be attributed to female preference for those males. However, female preference for less bold males would disagree with previous observations of female choice in the species (see below) (Scherer et al., 2020; Scherer, Kuhnhardt, et al., 2017). Another possibility is that a high level of male boldness might be associated with high aggressiveness (Sih et al., 2004; Tamin et al., 2023) leading to bolder males displaying increased aggressive behaviour towards females, which, in turn, could result in fewer egg laying or fertilization events.

Although male boldness expressed before breeding was associated with reproductive success, there was no link between female pre-breeding boldness and reproductive success. Previous mate choice studies on the rainbow krib yielded a similar pattern: female boldness did not predict male mate choice (Scherer & Schuett, 2018) but male boldness mattered during female mate choice (Scherer et al., 2020; Scherer, Kuhnhardt, et al., 2017). More specifically, in a correlative mate choice experiment, females preferred males of a dis-similar level of boldness (activity under simulated predation risk) to themselves (Scherer, Kuhnhardt, et al., 2017). And in a manipulative mate choice experiment (where boldness was manipulated using a gradient in ambient water temperature), indirect evidence for female preference for bold over less bold males was found: when given the choice between two males, female preference for the apparently bold male increased with increasing contrast in apparent male boldness between the two males presented (Scherer et al., 2020). We acknowledge there is some uncertainty regarding the direction of effect of those two mate choice studies, potentially because males presented in the manipulative mate choice study were generally bolder than females; in such a constellation, it is not necessarily possible to distinguish between a directional preference for bold males and a preference for dis-similar males (Scherer et al., 2020). However, by demonstrating male but not female mate choice for boldness in the rainbow krib, these previous findings further strengthen the prediction that male boldness may covary with a male-specific trait, such as sperm quality (see above).

Individual differences in boldness assessed before breeding did not predict individual differences in parental care behaviour, indicating that individuals might adjust their behaviour in response to the breeding conditions. Most importantly, breeding conditions may vary among breeding pairs with respect to characteristics of the social environment, including brood size (Sahm et al., 2023), offspring quality (Thünken et al., 2010), partner quality (Robart & Sinervo, 2019), and partner behaviour (Westneat et al., 2011). Individuals having to adapt to these different conditions during the breeding might have caused the lack of cross-context correlation of behaviour. Additionally, the way individuals adjust their behaviour to the breeding environment may also differ between behavioural types (Dingemanse et al., 2010; Laubu et al., 2016), which could have further decreased the predictive power of behaviour shown before breeding on behaviour shown during breeding. Alternatively, even though we selected a behaviour seemingly relevant for parental care, i.e. boldness towards a predator, which could be used as indicator of the ability to defend offspring, other behaviours, that we did not assess might be more important predictors for parental care, e.g. aggression.

Females that showed higher parental boldness had more offspring. In the literature, cichlids are often described as showing traditional role allocation with males engaging more in indirect care and females being the primary direct-care givers (Lavery & Reebs, 2010; McKaye & Murry, 2008; Richter et al., 2005). However, our result indicates that also female indirect care might be of paramount importance though future studies will need to investigate the causality of this correlation: do females that have more offspring (or a high-quality partner) become more invested (see above) or do females that show high parental boldness have an inherent quality that causes both, high parental boldness as well as many offspring?

Taken together, male but not female boldness predicted reproductive success though the link was not mediated via behavioural benefits during parental care, indicating a potential link to male genetic quality. Our results further indicate male boldness might be associated with different reproductive strategies (bold males sired fewer but bigger broods while less bold males sired more but smaller broods) that have the potential to maintain among-individual variation in male boldness if they lead to equal average fitness (e.g. caused by environmental heterogeneity). Interestingly, female parental boldness was correlated with the number of offspring produced highlighting the potential importance of female indirect care, however, the causality of this connection needs further investigation. In conclusion, our work shows that male boldness has important, complex fitness implications and highlights the need to unravel the role of genetic influences and environmental dynamics as potential key players shaping personality variation.

## Acknowledgements

We thank Milena Markwart for assistance in animal maintenance. Our research was funded by DFG (Deutsche Forschungsgemeinschaft, grant to WS: SCHU-2927/2-1).

## Notes

### Competing Interest Statement

The authors have declared no competing interest.

