## Supplementary information for "Male boldness and female parental care predict reproductive success in a bi-parental cichlid, the rainbow krib (*Pelvicachromis pulcher*)"

SUPPLEMENTARY INFORMATION 3

1 INTRUDER TYPE DID NOT AFFECT PARENTAL CARE BEHAVIOUR 3

2 FULL AND FINAL MODEL SUMMARIES 7

3 BEHAVIOURAL PROFILES OF MALES AND FEMALES USED IN THE BREEDING EXPERIMENT 13

### SUPPLEMENTARY INFORMATION

#### 1 Intruder type did not affect parental care behaviour

To assess whether the intruder type influenced parental care behaviour, we built two linear mixed-effect models with either parental boldness or brood guarding as the response (N = 120, respectively). As predictors we included intruder type (male conspecific, female conspecific, predator) and boldness shown before breeding as well as the interaction term between pre-breeding boldness and intruder type to test for the possibility that behavioural types may differ in their intruder differentiation. We further included trial number (one to six) as covariate, and test fish ID as well as family as random terms. Models were run for males and females separately. We found no effect of intruder type on parental boldness (**Figure S1a**, **Table S1**) or brood guarding (**Figure S1b**, **Table S2**). Pre-breeding boldness did not predict intruder differentiation either in interaction with intruder type or as main effect (**Table S1** and **S2**).

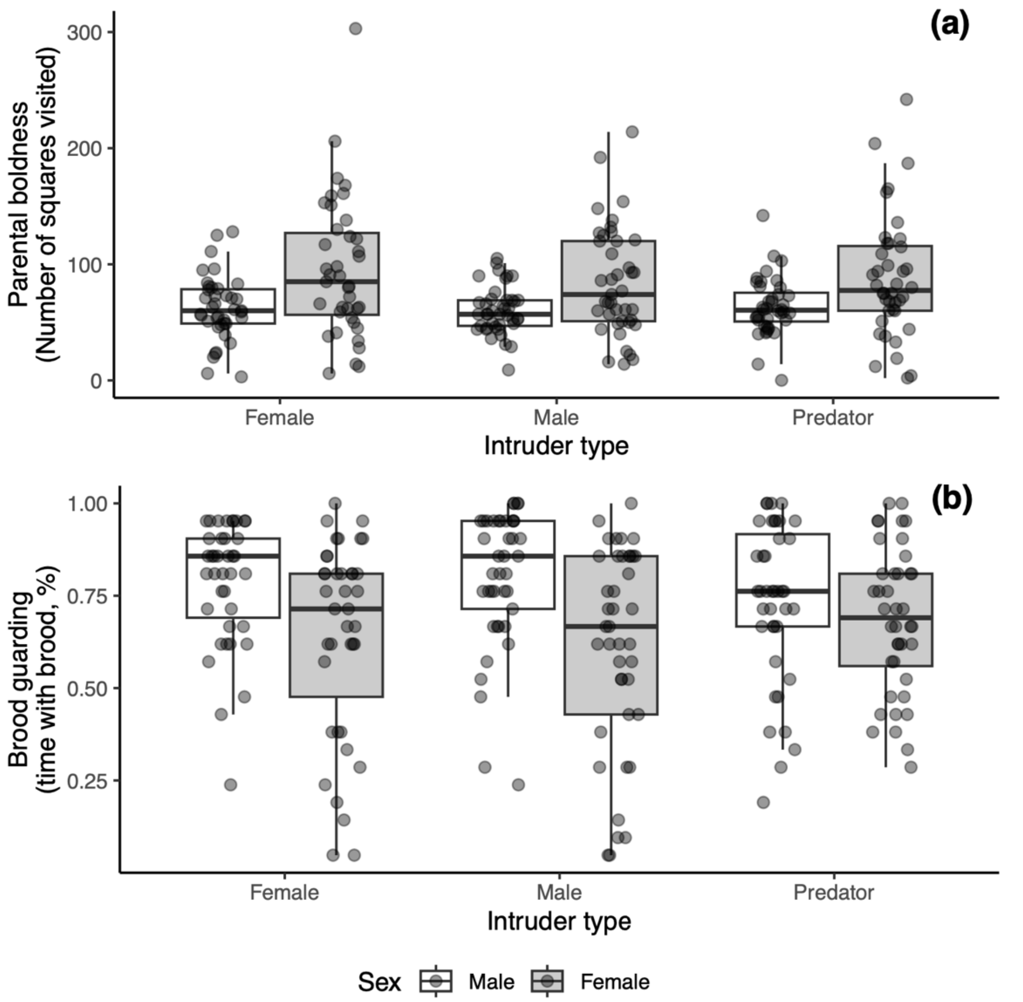


**Figure S1** No effect of intruder type on (a) parental boldness or (b) brood guarding in males (white boxes) or females (grey boxes). Shown are medians (horizontal lines), 25% and 75% quantiles (boxes), and 1.5 x interquartile ranges (whiskers).

**Table S1** LMMs testing if intruder type affects parental boldness, and if the response towards different intruder types depends on pre-breeding boldness. Results are controlled for trial number wherever necessary. Presented are full models (left, containing all predictors) and final models (right, containing significant predictors only), models were run for males (top) and females (bottom) separately. Boldness was assessed as the number of squares visited throughout.

| *Response* | *Predictors* | | Full model | | | Final model | | |
|---|---|---|---|---|---|---|---|---|
| | | | *Estimates* | *SE* | *p* | *Estimates* | *SE* | *p* |
| Parental boldness | Males | (Intercept) | 122.376 | 24.615 | - | 89.050 | 8.013 | - |
| | | Male pre-breeding boldness | -0.464 | 0.378 | - | - | - | - |
| | | Intruder type [male] | -43.080 | 22.799 | - | - | - | - |
| | | Intruder type [predator] | -15.335 | 22.807 | | - | - | |
| | | Parental care trial number | -0.619 | 2.228 | 0.781 | - | - | - |
| | | Male pre-breeding boldness × Intruder type [male] | 0.610 | 0.368 | 0.235 | - | - | - |
| | | Male pre-breeding boldness × Intruder type [predator] | 0.163 | 0.368 | | - | - | |
| | | **Random Effects** | | | | | | |
| | | **$\sigma^2$** | 1735.38 | | | 1801.76 | | |
| | | $\tau_{00}$ | 960.81 Male ID | | | 983.84 Male ID | | |
| | | | 0.00 Male family | | | 0.00 Male family | | |
| | | N | 20 Male ID | | | 20 Male ID | | |
| | | | 4 Male family | | | 4 Male family | | |
| | | Observations | 120 | | | 120 | | |
| | | Marginal $R^2$ / Conditional $R^2$ | 0.047 / NA | | | 0.000 / NA | | |
| | Females | (Intercept) | 55.652 | 8.704 | - | 49.726 | 4.967 | - |
| | | Female pre-breeding boldness | -0.178 | 0.212 | - | - | - | - |
| | | Intruder type [male] | -0.859 | 9.559 | - | - | - | - |
| | | Intruder type [predator] | 3.375 | 9.607 | | - | - | |
| | | Parental care trial number | 3.541 | 1.181 | 0.003 | 3.504 | 1.172 | 0.003 |
| | | Female pre-breeding boldness × Intruder type [male] | -0.041 | 0.266 | 0.954 | - | - | - |
| | | Female pre-breeding boldness × Intruder type [predator] | -0.082 | 0.269 | | - | - | |
| | | **Random Effects** | | | | | | |
| | | **$\sigma^2$** | 479.95 | | | 480.78 | | |
| | | $\tau_{00}$ | 52.64 Female ID | | | 71.78 Female ID | | |
| | | | 0.00 Female family | | | 0.00 Female family | | |
| | | N | 19 Female ID | | | 19 Female ID | | |
| | | | 2 Female family | | | 2 Female family | | |
| | | Observations | 120 | | | 120 | | |
| | | Marginal $R^2$ / Conditional $R^2$ | 0.103 / NA | | | 0.070 / NA | | |

**Table S2** LMM results testing if intruder type affects brood guarding, and if the response towards different intruder types depends on pre-breeding boldness. Results are controlled for trial number wherever necessary. Presented are full models (left, containing all predictors) and final models (right, containing significant predictors only), models were run for males (top) and females (bottom) separately. Boldness was assessed as the number of squares visited throughout.

| *Response* | *Predictors* | | Full model | | | Final model | | |
|---|---|---|---|---|---|---|---|---|
| | | | *Estimates* | *std. Error* | *p* | *Estimates* | *std. Error* | *p* |
| Brood guarding | Males | (Intercept) | 0.353 | 0.105 | - | 0.547 | 0.051 | - |
| | | Male pre-breeding boldness | 0.003 | 0.002 | - | - | - | - |
| | | Intruder type [male] | 0.060 | 0.118 | - | - | - | - |
| | | Intruder type [predator] | 0.159 | 0.118 | | - | - | |
| | | Parental care trial number | 0.028 | 0.012 | 0.017 | 0.028 | 0.012 | 0.018 |
| | | Male pre-breeding boldness × Intruder type [male] | -0.002 | 0.002 | 0.527 | - | - | - |
| | | Male pre-breeding boldness × Intruder type [predator] | -0.002 | 0.002 | | - | - | |
| | | **Random Effects** | | | | | | |
| | | **σ**$^2$ | 0.05 | | | 0.05 | | |
| | | $\tau_{00}$ | 0.01 Male ID | | | 0.01 Male ID | | |
| | | | 0.00 Male family | | | 0.00 Male family | | |
| | | N | 20 Male ID | | | 20 Male ID | | |
| | | | 4 Male family | | | 4 Male family | | |
| | | Observations | 120 | | | 120 | | |
| | | Marginal R2 / Conditional R2 | 0.126 / NA | | | 0.046 / NA | | |
| | Females | (Intercept) | 0.706 | 0.071 | - | 0.688 | 0.040 | - |
| | | Female pre-breeding boldness | -0.000 | 0.002 | - | - | - | - |
| | | Intruder type [male] | -0.054 | 0.071 | - | - | - | - |
| | | Intruder type [predator] | -0.064 | 0.071 | | - | - | |
| | | Parental care trial number | 0.026 | 0.009 | 0.004 | 0.025 | 0.009 | 0.007 |
| | | Female pre-breeding boldness × Intruder type [male] | 0.002 | 0.002 | 0.477 | - | - | - |
| | | Female pre-breeding boldness × Intruder type [predator] | 0.000 | 0.002 | | - | - | |
| | | **Random Effects** | | | | | | |
| | | **σ**$^2$ | 0.03 | | | 0.03 | | |
| | | $\tau_{00}$ | 0.01 Female ID | | | 0.01 Female ID | | |
| | | | 0.00 Female family | | | 0.00 Female family | | |
| | | N | 19 Female ID | | | 19 Female ID | | |
| | | | 2 Female family | | | 2 Female family | | |
| | | Observations | 120 | | | 120 | | |
| | | Marginal $R^2$ / Conditional $R^2$ | 0.107 / NA | | | 0.060 / NA | | |

#### 2 Full and final model summaries

**Table S3** LMM results testing if average pre-breeding boldness (male behaviour, female behaviour, and the combination of male-female behaviour) predict a pair's likelihood to reproduce. Presented are full models (left, containing all predictors) and final models (right, containing significant predictors only). Boldness was assessed as the number of squares visited throughout.

| *Response* | *Predictors* | Full model | | | Final model | | |
|---|---|---|---|---|---|---|---|
| | | *Odds Ratios* | *std. Error* | *p* | *Odds Ratios* | *std. Error* | *p* |
| Likelihood to reproduce (yes/no) | (Intercept) | 23.328 | 47.689 | - | 1.882 | 2.214 | - |
| | Male pre-breeding boldness | 0.958 | 0.022 | - | 0.978 | 0.012 | 0.049 |
| | Female pre-breeding boldness | 0.944 | 0.040 | - | - | - | - |
| | Male pre-breeding boldness × Female pre-breeding boldness | 1.000 | 0.001 | 0.375 | - | - | - |
| | **Random Effects** | | | | | | |
| | $\sigma^2$ | 3.29 | | | 3.29 | | |
| | $\tau_{00}$ | 0.40 Male family | | | 0.62 Male family | | |
| | | 0.55 Female family | | | 1.28 Female family | | |
| | ICC | 0.22 | | | 0.37 | | |
| | N | 4 Male family | | | 4 Male family | | |
| | | 2 Female family | | | 2 Female family | | |
| | Observations | 54 | | | 54 | | |
| | Marginal $R^2$ / Conditional $R^2$ | 0.176 / 0.360 | | | 0.086 / 0.421 | | |
| | AIC | 70.110 | | | 68.762 | | |
| | log-Likelihood | -29.055 | | | -30.381 | | |

**Table S4** LMM results testing if pre-breeding boldness (male behaviour, female behaviour, and the combination of male-female behaviour) predict a pair's reproductive success regarding the number (top) and size (bottom) of offspring produced. Presented are full models (left, containing all predictors) and final models (right, containing significant predictors only). Boldness was assessed as the number of squares visited throughout.

| *Response* | *Predictors* | Full model | | | Final model | | |
|---|---|---|---|---|---|---|---|
| | | *Estimates* | *std. Error* | *p* | *Estimates* | *std. Error* | *p* |
| Number of offspring | (Intercept) | 45.413 | 40.042 | - | 29.618 | 27.260 | - |
| | Male pre-breeding boldness | 0.635 | 0.414 | - | 0.800 | 0.193 | 0.001 |
| | Female pre-breeding boldness | -0.542 | 1.084 | - | - | - | - |
| | Male pre-breeding boldness × Female pre-breeding boldness | 0.007 | 0.016 | 0.690 | - | - | - |
| | **Random Effects** | | | | | | |
| | $\sigma^2$ | 377.16 | | | 385.53 | | |
| | $\tau_{00}$ | 2440.61 Male family | | | 2251.27 Male family | | |
| | | 57.45 Female family | | | 86.83 Female family | | |
| | ICC | 0.87 | | | 0.86 | | |
| | N | 2 Female family | | | 2 Female family | | |
| | | 4 Male family | | | 4 Male family | | |
| | Observations | 20 | | | 20 | | |
| | Marginal R2 / Conditional R2 | 0.139 / 0.887 | | | 0.137 / 0.878 | | |
| Average offspring size (cm) | (Intercept) | 1.582 | 0.153 | - | 1.567 | 0.026 | - |
| | Male pre-breeding boldness | -0.001 | 0.002 | - | - | - | - |
| | Female pre-breeding boldness | -0.001 | 0.005 | - | - | - | - |
| | Male pre-breeding boldness × Female pre-breeding boldness | 0.000 | 0.000 | 0.667 | - | - | - |
| | **Random Effects** | | | | | | |
| | $\sigma^2$ | 0.01 | | | 0.01 | | |
| | $\tau_{00}$ | 0.00 Male family | | | 0.00 Male family | | |
| | | 0.00 Female family | | | 0.00 Female family | | |
| | N | 2 Female family | | | 2 Female family | | |
| | | 4 Male family | | | 4 Male family | | |
| | Observations | 20 | | | 20 | | |
| | Marginal $R^2$ / Conditional $R^2$ | 0.028 / NA | | | 0.000 / NA | | |

**Table S5** LMM results testing if pre-breeding boldness predicts parental boldness; LMMs were run for male (top) and female (middle) behaviour as well as the combination of thereof (aka behavioural contrast; bottom). Presented are full models (left, containing all predictors) and final models (right, containing significant predictors only). Boldness was assessed as the number of squares visited throughout.

| *Response* | *Predictors* | Full model | | | Final model | | |
|---|---|---|---|---|---|---|---|
| | | *Estimates* | *std. Error* | *p* | *Estimates* | *std. Error* | *p* |
| Male parental boldness | (Intercept) | 100.691 | 19.395 | - | 89.050 | 8.013 | - |
| | Male pre-breeding boldness | -0.206 | 0.313 | 0.513 | - | - | - |
| | **Random Effects** | | | | | | |
| | $\sigma^2$ | 1256.95 | | | 1284.13 | | |
| | $\tau_{00}$ | 0.00 Male family | | | 0.00 Male family | | |
| | N | 4 Male family | | | 4 Male family | | |
| | Observations | 20 | | | 20 | | |
| | Marginal $R^2$ / Conditional $R^2$ | 0.022 / NA | | | 0.000 / NA | | |
| Female parental boldness | (Intercept) | 68.702 | 5.273 | - | 61.842 | 2.857 | - |
| | Female pre-breeding boldness | -0.221 | 0.146 | 0.140 | - | - | - |
| | **Random Effects** | | | | | | |
| | $\sigma^2$ | 146.39 | | | 163.20 | | |
| | $\tau_{00}$ | 0.00 Female family | | | 0.00 Female family | | |
| | N | 2 Female family | | | 2 Female family | | |
| | Observations | 20 | | | 20 | | |
| | Marginal $R^2$ / Conditional $R^2$ | 0.108 / NA | | | 0.000 / NA | | |
| Male-female contrast in parental boldness | (Intercept) | 31.850 | 10.269 | - | 27.208 | 8.378 | - |
| | Male-female contrast in pre-breeding boldness | -0.182 | 0.239 | 0.450 | - | - | - |
| | **Random Effects** | | | | | | |
| | $\sigma^2$ | 1364.34 | | | 1403.81 | | |
| | $\tau_{00}$ | 0.00 Male family | | | 0.00 Male family | | |
| | | 0.00 Female family | | | 0.00 Female family | | |
| | N | 2 Female family | | | 2 Female family | | |
| | | 4 Male family | | | 4 Male family | | |
| | Observations | 20 | | | 20 | | |
| | Marginal $R^2$ / Conditional $R^2$ | 0.030 / NA | | | 0.000 / NA | | |

**Table S6** LMM results testing if pre-breeding boldness predicts brood guarding. Presented are full models (left, containing all predictors) and final models (right, containing significant predictors only), models were run for male (top) and female (middle) behaviour as well as the combination of thereof (aka behavioural contrast; bottom). Boldness was assessed as the number of squares visited throughout.

| *Response* | *Predictors* | **Full model** | | | **Final model** | | |
|---|---|---|---|---|---|---|---|
| | | *Estimates* | *std. Error* | *p* | *Estimates* | *std. Error* | *p* |
| Male brood guarding | (Intercept) | 0.524 | 0.069 | - | 0.646 | 0.031 | - |
| | Male pre-breeding boldness | 0.002 | 0.001 | 0.063 | - | - | - |
| | **Random Effects** | | | | | | |
| | $\sigma^2$ | 0.02 | | | 0.02 | | |
| | $\tau_{00}$ | 0.00 Male family | | | 0.00 Male family | | |
| | N | 4 Male family | | | 4 Male family | | |
| | Observations | 20 | | | 20 | | |
| | Marginal $R^2$ / Conditional $R^2$ | 0.166 / NA | | | 0.000 / NA | | |
| Female brood guarding | (Intercept) | 0.756 | 0.046 | - | 0.774 | 0.024 | - |
| | Female pre-breeding boldness | 0.001 | 0.001 | 0.656 | - | - | - |
| | **Random Effects** | | | | | | |
| | $\sigma^2$ | 0.01 | | | 0.01 | | |
| | $\tau_{00}$ | 0.00 Female family | | | 0.00 Female family | | |
| | N | 2 Female family | | | 2 Female family | | |
| | Observations | 20 | | | 20 | | |
| | Marginal $R^2$ / Conditional $R^2$ | 0.010 / NA | | | 0.000 / NA | | |
| Male-female contrast in brood guarding | (Intercept) | -0.149 | 0.043 | - | -0.123 | 0.033 | - |
| | Male-female contrast in pre-breeding boldness | 0.002 | 0.001 | 0.066 | - | - | - |
| | **Random Effects** | | | | | | |
| | $\sigma^2$ | 0.01 | | | 0.02 | | |
| | $\tau_{00}$ | 0.00 Male family | | | 0.00 Male family | | |
| | | 0.00 Female family | | | 0.00 Female family | | |
| | N | 2 Female family | | | 2 Female family | | |
| | | 4 Male family | | | 4 Male family | | |
| | Observations | 20 | | | 20 | | |
| | Marginal $R^2$ / Conditional $R^2$ | 0.209 / NA | | | 0.000 / NA | | |

**Table S7** LMM results testing if parental boldness (top) or brood guarding (bottom) predict the number of offspring produced. We considered male behaviour, female behaviour, and the combination of thereof. Presented are full models (left, containing all predictors) and final models (right, containing significant predictors only). Boldness was assessed as the number of squares visited throughout.

| *Response* | *Predictors* | Full model | | | Final model | | |
|---|---|---|---|---|---|---|---|
| | | *Estimates* | *std. Error* | *p* | *Estimates* | *std. Error* | *p* |
| Number of offspring | (Intercept) | -43.000 | 78.410 | - | 7.083 | 35.131 | - |
| | Male parental boldness | 0.510 | 0.766 | - | - | - | - |
| | Female parental boldness | 2.028 | 1.375 | - | 1.029 | 0.465 | 0.041 |
| | Male parental boldness × Female parental boldness | -0.011 | 0.014 | 0.498 | - | - | - |
| | **Random Effects** | | | | | | |
| | $\sigma^2$ | 646.51 | | | 663.27 | | |
| | $\tau_{00}$ | 964.58 Male family | | | 1132.18 Male family | | |
| | | 114.10 Female family | | | 57.67 Female family | | |
| | ICC | 0.63 | | | 0.64 | | |
| | N | 2 Female family | | | 2 Female family | | |
| | | 4 Male family | | | 4 Male family | | |
| | Observations | 20 | | | 20 | | |
| | Marginal $R^2$ / Conditional $R^2$ | 0.115 / 0.668 | | | 0.089 / 0.674 | | |
| | (Intercept) | 103.136 | 342.282 | - | 74.465 | 18.887 | - |
| | Male brood guarding | -95.172 | 531.114 | - | - | - | - |
| | Female brood guarding | -97.164 | 455.513 | - | - | - | - |
| | Male brood guarding x Female brood guarding | 214.240 | 693.142 | 0.761 | - | - | - |
| | **Random Effects** | | | | | | |
| | $\sigma^2$ | 725.82 | | | 871.22 | | |
| | $\tau_{00}$ | 1108.17 Male family | | | 1210.45 Male family | | |
| | | 0.00 Female family | | | 0.00 Female family | | |
| | ICC | | | | 0.58 | | |
| | N | 2 Female family | | | 2 Female family | | |
| | | 4 Male family | | | 4 Male family | | |
| | Observations | 20 | | | 20 | | |
| | Marginal $R^2$ / Conditional $R^2$ | 0.177 / NA | | | 0.000 / 0.581 | | |

**Table S8** LMM results testing if parental boldness (top) or brood guarding (bottom) predict the size of offspring produced. We considered male behaviour, female behaviour, and the combination of thereof. Presented are full models (left, containing all predictors) and final models (right, containing significant predictors only). Boldness was assessed as the number of squares visited throughout.

| *Response* | *Predictors* | **Full model** | | | **Final model** | | |
|---|---|---|---|---|---|---|---|
| | | *Estimates* | *std. Error* | *p* | *Estimates* | *std. Error* | *p* |
| Average offspring size (cm) | (Intercept) | 1.851 | 0.293 | - | 1.567 | 0.026 | - |
| | Male parental boldness | -0.004 | 0.003 | - | - | - | - |
| | Female parental boldness | -0.006 | 0.005 | - | - | - | - |
| | Male parental boldness × Female parental boldness | 0.000 | 0.000 | 0.189 | - | - | - |
| | **Random Effects** | | | | | | |
| | $\sigma^2$ | 0.01 | | | 0.01 | | |
| | $\tau_{00}$ | 0.00 Male family | | | 0.00 Male family | | |
| | | 0.00 Female family | | | 0.00 Female family | | |
| | N | 2 Female family | | | 2 Female family | | |
| | | 4 Male family | | | 4 Male family | | |
| | Observations | 20 | | | 20 | | |
| | Marginal $R^2$ / Conditional $R^2$ | 0.099 / NA | | | 0.000 / NA | | |
| | (Intercept) | 1.582 | 1.317 | - | 1.567 | 0.026 | - |
| | Male brood guarding | 0.212 | 2.049 | - | - | - | - |
| | Female brood guarding | -0.034 | 1.734 | - | - | - | - |
| | Male brood guarding × Female brood guarding | -0.247 | 2.651 | 0.926 | - | - | - |
| | **Random Effects** | | | | | | |
| | $\sigma^2$ | 0.01 | | | 0.01 | | |
| | $\tau_{00}$ | 0.00 Male family | | | 0.00 Male family | | |
| | | 0.00 Female family | | | 0.00 Female family | | |
| | N | 2 Female family | | | 2 Female family | | |
| | | 4 Male family | | | 4 Male family | | |
| | Observations | 20 | | | 20 | | |
| | Marginal $R^2$ / Conditional $R^2$ | 0.031 / NA | | | 0.000 / NA | | |

#### 3 Behavioural profiles of males and females used in the breeding experiment

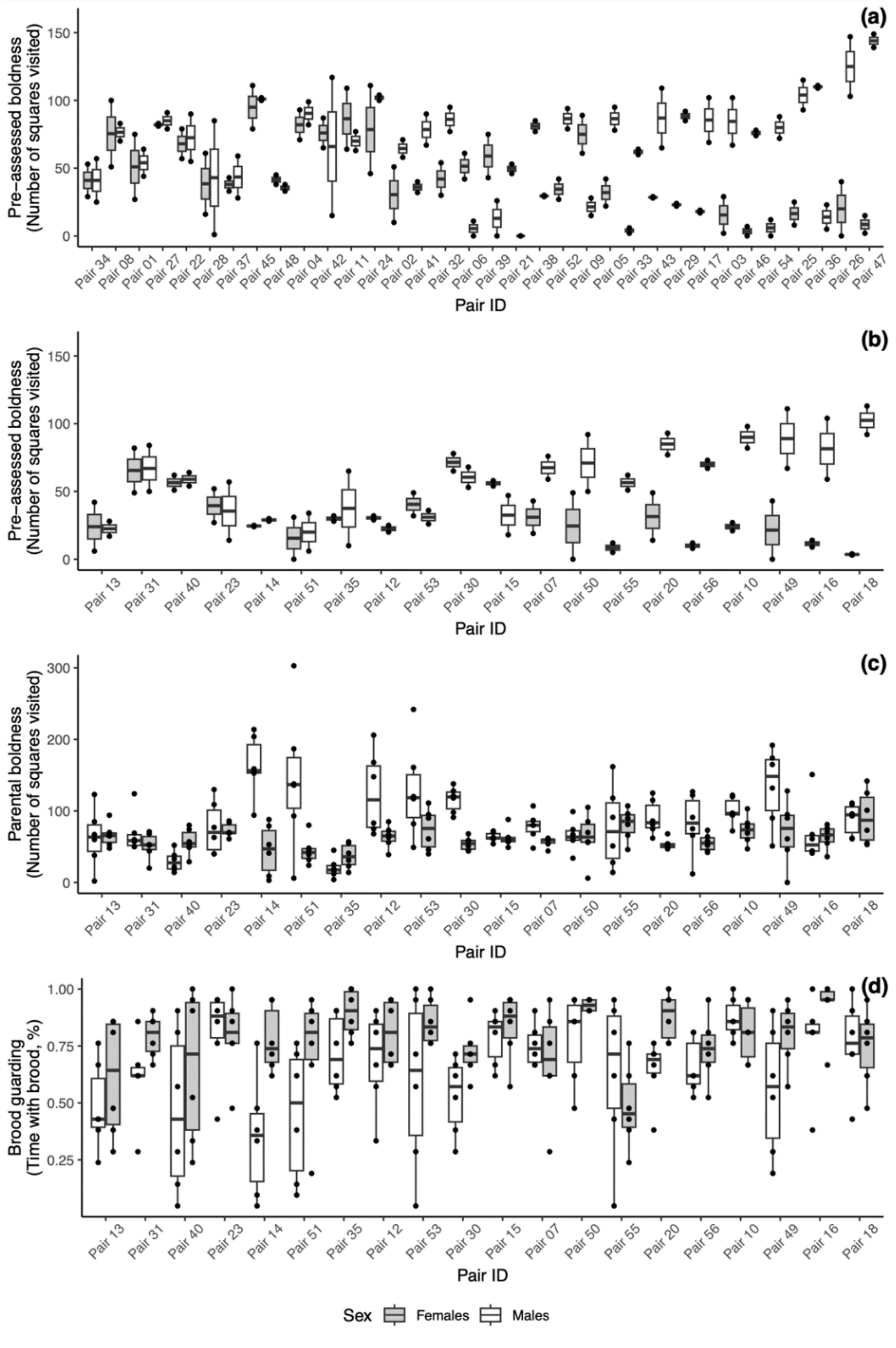


**Figure S2** Behavioural scores for all male and female rainbow kribs used in the breeding experiment. (a-b) Pre-breeding boldness scores of (a) unsuccessful vs. (b) successful pairs (two tests per ID) and (c-d) parental care behaviour (six tests per ID); all panels ordered by within-pair behavioural contrast in pre-breeding boldness (from smallest to largest contrast).
